# A C-terminal motif containing a PKC phosphorylation site regulates γ-Protocadherin-mediated dendrite arborization in the cerebral cortex *in vivo*

**DOI:** 10.1101/2024.01.25.577214

**Authors:** Camille M. Hanes, Kar Men Mah, David M. Steffen, Charles G. Marcucci, Leah C. Fuller, Robert W. Burgess, Andrew M. Garrett, Joshua A. Weiner

## Abstract

The *Pcdhg* gene cluster encodes 22 γ-Protocadherin (γ-Pcdh) cell adhesion molecules that critically regulate multiple aspects of neural development, including neuronal survival, dendritic and axonal arborization, and synapse formation and maturation. Each γ-Pcdh isoform has unique protein domains–a homophilically-interacting extracellular domain and a juxtamembrane cytoplasmic domain– as well as a C-terminal cytoplasmic domain shared by all isoforms. The extent to which isoform-specific *vs.* shared domains regulate distinct γ-Pcdh functions remains incompletely understood. Our previous *in vitro* studies identified PKC phosphorylation of a serine residue within a shared C-terminal motif as a mechanism through which γ-Pcdh promotion of dendrite arborization *via* MARCKS is abrogated. Here, we used CRISPR/Cas9 genome editing to generate two new mouse lines expressing only non-phosphorylatable γ-Pcdhs, due either to a serine-to-alanine mutation (*Pcdhg^S/A^*) or to a 15-amino acid C-terminal deletion resulting from insertion of an early stop codon (*Pcdhg^CTD^*). Both lines are viable and fertile, and the density and maturation of dendritic spines remains unchanged in both *Pcdhg^S/A^* and *Pcdhg^CTD^*cortex. Dendrite arborization of cortical pyramidal neurons, however, is significantly increased in both lines, as are levels of active MARCKS. Intriguingly, despite having significantly reduced levels of γ-Pcdh proteins, the *Pcdhg^CTD^* mutation yields the strongest phenotype, with even heterozygous mutants exhibiting increased arborization. The present study confirms that phosphorylation of a shared C-terminal motif is a key γ-Pcdh negative regulation point, and contributes to a converging understanding of γ-Pcdh family function in which distinct roles are played by both individual isoforms and discrete protein domains.

## INTRODUCTION

The development of the mammalian central nervous system is a tightly regulated process encompassing multiple overlapping stages, such as neurogenesis, neuronal migration, axon outgrowth, dendrite arborization, and synaptogenesis. Proper neural circuit formation during development is critical for later mature brain function, and many neurodevelopmental disorders have been associated with defective neural circuitry (reviewed in (Exposito-Alonso & Rico, 2022; Sanes & Zipursky, 2020)). Cell-cell interactions are critical to CNS development: both axons and dendrites rely on diverse cell adhesion molecule (CAM) interactions as they grow out, branch, and find appropriate synaptic partners. The clustered protocadherin (cPcdh) genes within the tandemly arrayed *Pcdha, Pcdhb,* and *Pcdhg* clusters encode a combined total of ∼60 distinct CAMs that provide molecular diversity to the cell surfaces of individual neurons (Canzio & Maniatis, 2019; Wu & Maniatis, 1999; Wu et al., 2001). In the mouse, the *Pcdha* cluster encodes 12 highly similar isoforms (α-Pcdhs 1-12) and two more distantly related isoforms, αC1 and αC2, the *Pcdhb* cluster encodes 22 isoforms, and the *Pcdhg* cluster encodes 19 highly similar isoforms (12 γ-Pcdh-A subtype proteins and 7 γ-Pcdh-B subtype proteins) as well as 3 that are most highly related to the α-Pcdh C-type isoforms (γC3, γC4, and γC5; Fig. 1A). Each α-Pcdh and γ-Pcdh protein is encoded by four exons. The first large “variable” exon encodes 6 extracellular cadherin (EC) repeats, one transmembrane domain, and a ∼90 amino acid variable cytoplasmic domain (VCD). Each variable exon is spliced to three short “constant” exons, which together encode a 125-amino acid shared C-terminal domain. In contrast, the β-Pcdhs lack any such shared constant domain and are each encoded by a single large variable exon (Wu et al., 2001)(Fig. 1A).

**Figure 1.**
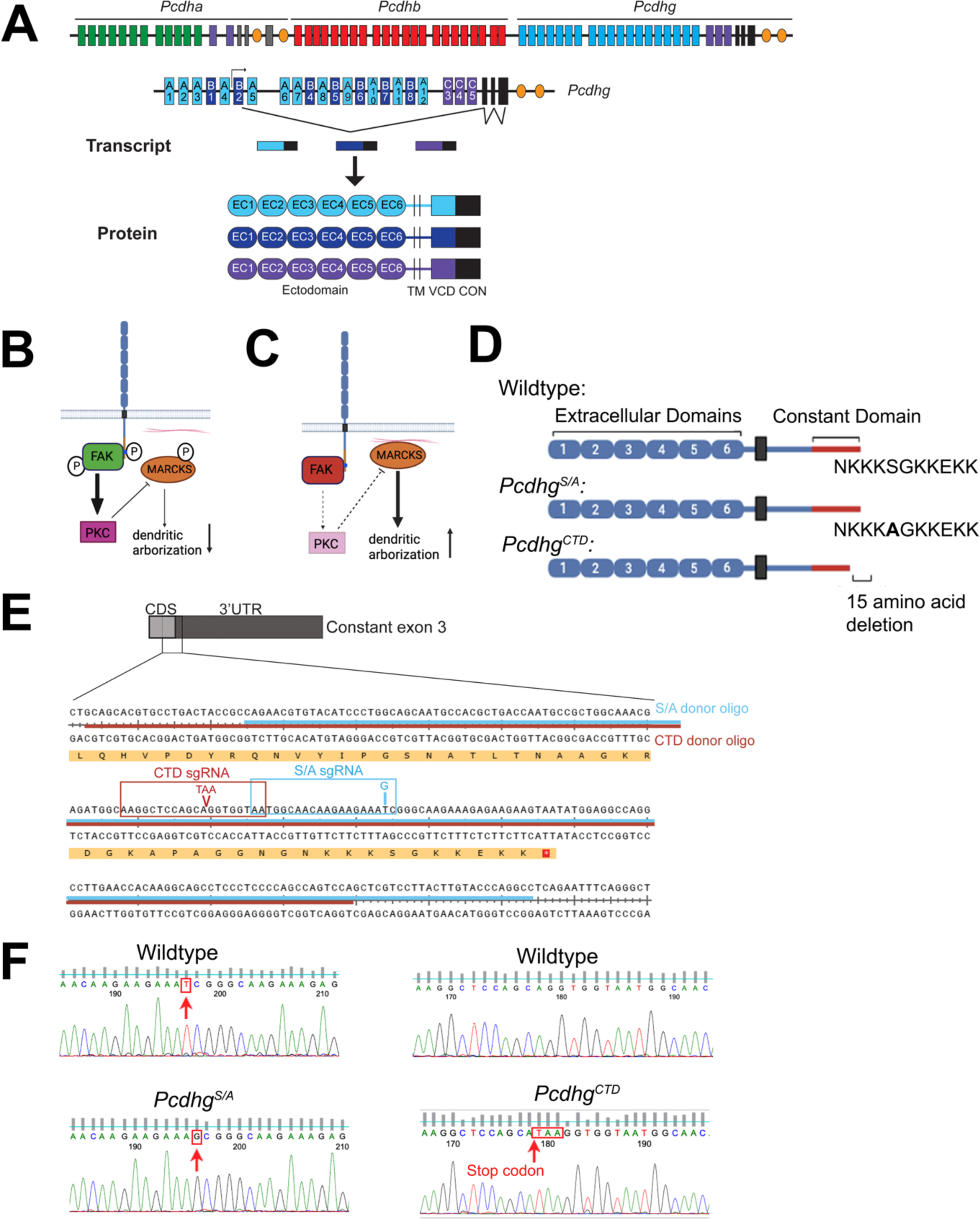
Generation of the *Pcdhg^S/A^*and *Pcdhg^CTD^* mice. **A**, Schematic representation of (top to bottom): the mouse clustered protocadherin loci, comprising *Pcdha*, *Pcdhb*, and *Pcdhg* clusters; the *Pcdhg* locus, comprising 22 variable exons (γA-subtype: light blue, γB-subtype: dark blue, and γC-subtype: purple) which are spliced to three constant exons (black); *Pcdhg* transcripts comprising one large variable exon and three constant exons; and the resultant γ-Pcdh protein domain structure, with six extracellular cadherin (EC) repeats, a transmembrane (TM) domain, a short variable cytoplasmic domain (VCD; all encoded by the variable exon) followed by a shared cytoplasmic domain (encoded by the three constant exons). **B**, Schematic of the FAK/PKC/MARCKS signaling pathway identified in Garrett et al. (2012) and Keeler et al. (2015) in the presence (**B**) and absence (**C**) of γ-Pcdh constant domain phosphorylation. Pink lines represent actin filaments. **D**, Schematic representation of (top to bottom): the wildtype γ-Pcdh protein with an intact constant domain; the serine-to-alanine mutation in *Pcdhg^S/A^*, and the C-terminal deletion in *Pcdhg^CTD^*. **E**, sgRNAs targeting the third constant exon to generate *Pcdhg^S/A^* (blue) and *Pcdhg^CTD^* (red), resulting in a thymine-to-guanine point mutation and insertion of an early stop codon, respectively. **F**, Chromatograms of *Pcdhg^S/A^* and *Pcdhg^CTD^* genomic DNA sequencing compared to wildtype.

Over 20 years of functional studies have identified the γ-Pcdhs as the only cPcdhs essential for neonatal viability, and they additionally play the most important roles in several key steps in neural development (reviewed in (Peek et al., 2017)). Excessive spinal interneuron apoptosis preceding neonatal lethality in mice lacking all 22 *Pcdhg* genes indicates the importance of these molecules in organismal survival (Lefebvre et al., 2008; Mancia Leon et al., 2024; Prasad et al., 2008; Wang et al., 2002). In the CNS, loss of all γ-Pcdhs or disruption of normal γ-Pcdh repertoire consistently leads to disrupted neurodevelopment, including reduced dendritic arborization, loss of dendrite and axon self-avoidance, aberrant axon coalescence, and/or altered synapse formation/maturation depending upon the neuronal cell type (Garrett et al., 2012; Garrett & Weiner, 2009; Ing-Esteves et al., 2018; Kobayashi et al., 2023; Kostadinov & Sanes, 2015; Lefebvre et al., 2012; Molumby et al., 2017; Molumby et al., 2016; Mountoufaris et al., 2017; Prasad & Weiner, 2011; Steffen et al., 2021; Weiner et al., 2005).

While initial analyses utilized mouse models in which the entire *Pcdhg* locus was disrupted, more recent work has made progress in identifying isoform-specific roles among the 22 distinct γ-Pcdhs, particularly for the three C-type isoforms. A shotgun CRISPR/Cas9-mediated mutagenesis screen of the *Pcdhg* variable exons (Garrett et al., 2019) resulted in the surprising demonstration that the γC4 isoform is the only one required for organismal survival: mice carrying a discrete loss-of-function mutation in γC4 died shortly after birth with excessive spinal interneuron apoptosis, despite expression of the other 21 γ-Pcdhs. Conversely, a mutant mouse line in which γC4 is the only remaining functional isoform is viable and fertile (Garrett et al., 2019). This unique role for γC4 in neuronal survival has recently been confirmed for interneurons in the developing neocortex as well (Mancia Leon et al., 2024). We found that the γC3 VCD uniquely inhibits canonical Wnt signaling in non-neuronal cells through its direct interaction with the scaffolding protein Axin1 (Mah et al., 2016). Consistent with this, discrete loss of the γC3 isoform in mice results in a reduction in cortical dendrite arborization nearly as great as that observed when all γ-Pcdhs are lost, and this can be rescued by re-expression of γC3 in an Axin1-dependent manner (Steffen et al., 2023). Additionally, restricted overexpression of γC3 in the cortex was reported to affect the lateral dispersion of clonally-related neurons, disrupting the preferential pattern of synaptic connectivity between them (Lv et al., 2022). A further role for γC3 has been discovered in the spinal cord, where it preferentially acts to promote proper sensory neuron synapse formation (Meltzer et al., 2023). Finally, γC5 has been reported to interact with the γ2 subunit of GABA_A_ receptors via their cytoplasmic domains, and this facilitates GABA_A_ receptor cell surface expression and GABAergic synapse stability and maintenance (Li et al., 2017; Li et al., 2012).

These proof-of-principle demonstrations of isoform-specific γ-Pcdh roles (albeit limited thus far to the C-type isoforms, which are the most highly and broadly expressed) raise the question of constant domain functions, which reflect all 22 γ-Pcdh proteins. A key insight into the role of the γ-Pcdh constant domain’s function was provided by Chen et al. (2009), who discovered a physical and functional interaction of the constant domain of γ-Pcdhs (as well as that of α-Pcdhs) with Focal Adhesion Kinase (FAK; also the FAK homologue Pyk2). The γ-Pcdh constant domain binds to FAK and inhibits its autophosphorylation (and thus activation) at Y397 (Chen et al., 2009). FAK is well-known to negatively regulate dendritic arborization (Armendariz et al., 2014; Rico et al., 2004), and consistent with this, dendritic arborization is severely decreased in the *Pcdhg* mutant cerebral cortex, while FAK activity is upregulated (Garrett et al., 2012). We subsequently elaborated a signaling pathway through which the γ-Pcdhs promoted cortical dendrite arborization (Garrett et al., 2012): downstream of FAK activation, protein kinase C (PKC) phosphorylates, and thus inhibits, myristoylated alanine-rich C-kinase substrate (MARCKS), an actin-binding protein crucial for normal arborization of dendrites (Li et al., 2008). By inhibiting FAK *via* their constant domains, γ-Pcdh proteins reduce PKC activity, resulting in MARCKS hypophosphorylation and allowing normal dendritic arbor complexity ((Garrett et al., 2012); Fig. 1C).

Keeler et al. (2015) subsequently showed that the γ-Pcdh constant domain C-terminus is, in turn, a substrate for phosphorylation by PKC at a particular serine residue (S119, numbered in reference to the 125 amino acids of the constant domain) embedded in a lysine-rich phospholipid-binding motif (Fig. 1D). Phosphorylation of S119 by PKC in HEK293 cells *in vitro* abrogates γ-Pcdh inhibition of (but not binding to) FAK, thus amplifying its downstream signaling pathways (Keeler et al., 2015; Fig. 1B). Consistent with this, transfecting cultured wildtype neurons with a γ-Pcdh construct harboring a serine-to-alanine mutation (Fig. 1D) that prevents constant domain phosphorylation by PKC (but not a wildtype γ-Pcdh construct) resulted in increased dendrite arborization (Keeler et al., 2015). Here, we sought to expand on these *in vitro* findings by testing the role of the shared γ-Pcdh C-terminus in the development of the cerebral cortex *in vivo.* We used CRISPR/Cas9 genome editing to generate two novel mouse lines in which the C-terminus of all 22 endogenous γ-Pcdhs are unable to be phosphorylated by PKC due either to a point mutation resulting in a serine-to-alanine substitution (S119A; *Pcdhg^S/A^*) or the insertion of a premature stop codon that results in loss of the final 15 amino acids, removing both the PKC target S119 and the phospholipid-binding, lysine-rich motif (*Pcdhg^CTD^*, for “C-terminal deletion”). We find that both mouse lines are viable and fertile, and that overall brain structure is normal, indicating that the critical function of the γ-Pcdh protein family in neuronal survival and synapse formation is not significantly affected by shared C-terminus disruption. Analysis of cortical dendrite arborization, however, confirms our identification of S119 phosphorylation as a key negative regulation point for the γ-Pcdhs, as both *Pcdhg^S/A^* and *Pcdhg^CTD^* homozygous mice exhibit significantly over-exuberant arbors and decreased MARCKS phosphorylation. These results delineate unique functional contributions of a particular γ-Pcdh protein domain and, along with other recent studies genetically dissecting the *Pcdhg* locus *in vivo* (Garrett et al., 2019; Mancia Leon et al., 2024; Meltzer et al., 2023; Molumby et al., 2017; Molumby et al., 2016; Steffen et al., 2021; Steffen et al., 2023), contribute to a better understanding of how this diverse protein family regulates the development of a functioning nervous system.

## RESULTS

### Generation of two novel γ-Pcdh constant domain mutant mouse line, *Pcdhg^S/A^* and *Pcdhg^CTD^*

To investigate the role of γ-Pcdh constant domain phosphorylation *in vivo*, we used CRISPR/Cas9 genome editing to generate two mutant mouse lines in which the constant domain cannot be phosphorylated. In the first, referred to as *Pcdhg^S/A^*, a thymine-to-guanine point mutation was introduced that results in a serine-to-alanine (S/A) mutation at amino acid residue 119 (of the constant domain’s 125), thus preventing phosphorylation at this site by PKC (Fig. 1D, E). In the second mouse line, referred to as *Pcdhg^CTD^*, the insertion of a premature stop codon (TAA) at the 3’ end of the third constant exon results in a 15 amino acid C-terminal deletion in all γ-Pcdhs that includes S119 and the phospholipid-binding motif in which it is situated (CTD; Fig. 1D, E). Genomic sequencing confirmed that the intended mutations were introduced (Fig. 1F). Both *Pcdhg^S/A^*and *Pcdhg^CTD^* homozygotes were viable and fertile, and gross brain structure was normal, with no apparent disruption to cerebral cortex layer composition, cell survival, or density of molecularly-defined cell types (Fig. 2A).

**Figure 2.**
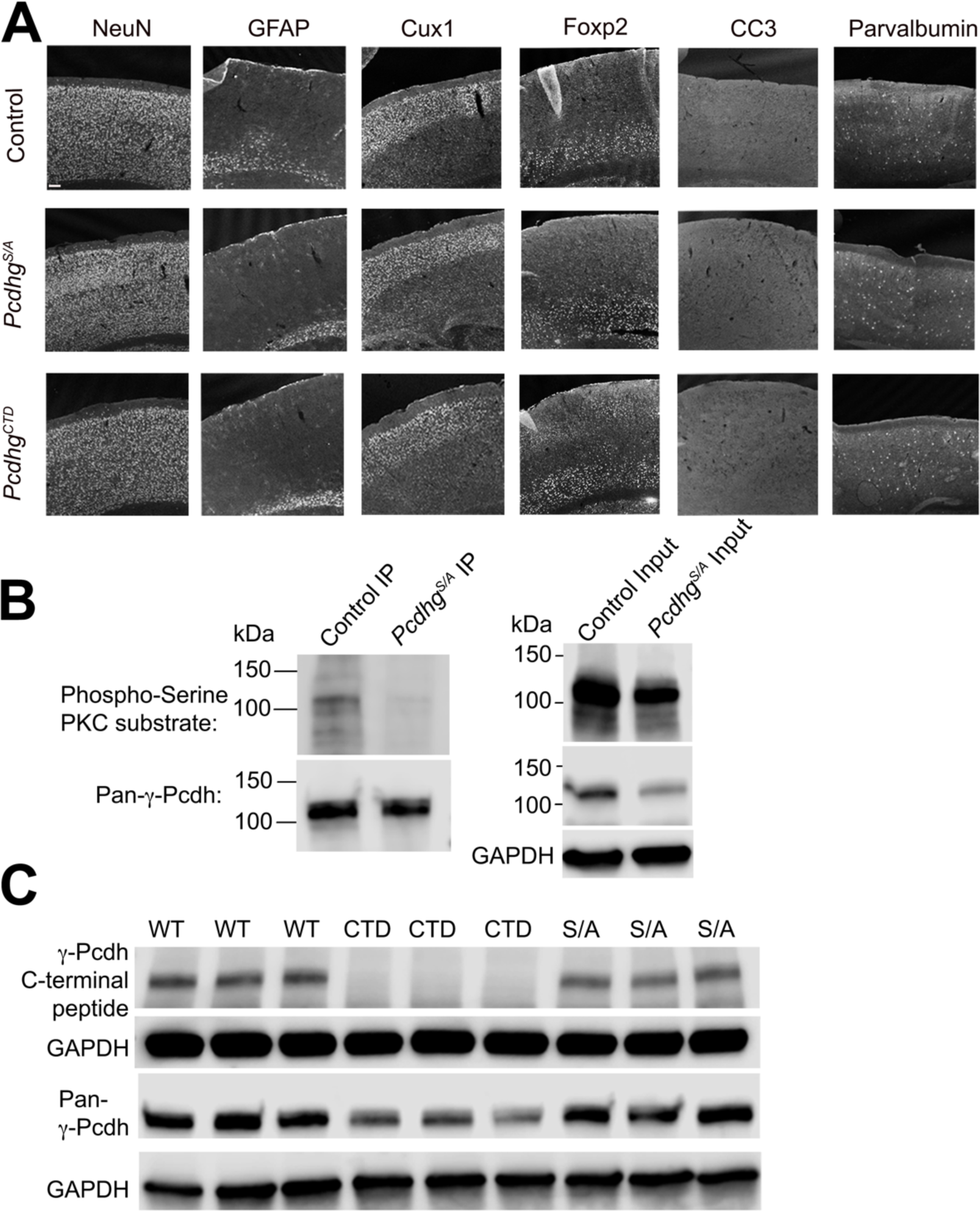
Validation of the *Pcdhg^S/A^* and *Pcdhg^CTD^* mice. **A**, Cryosections of adult control, *Pcdhg^S/A^*, and *Pcdhg^CTD^* homozygous mutant cortices stained with the indicated layer- and cell type-specific markers (NeuN, neurons; GFAP, astrocytes; Cux1, upper layer neurons; FoxP2, deep layer neurons; cleaved caspase-3 [CC3], apoptotic cells; Parvalbumin, interneuron subset), revealing grossly normal morphology, neuronal survival, and cell types in *Pcdhg^S/A^* and *Pcdhg^CTD^* mutants. Scale bar: 100 μm. **B**, Immunoprecipitation of γ-Pcdhs using a pan-γ-Pcdh antibody from control and *Pcdhg^S/A^* homozygous mutant brain lysates followed by blotting with an antibody specific to phosphoserine PKC substrates, revealing loss of PKC-phosphorylated γ-Pcdhs in *Pcdhg^S/A^*. **C**, Western blot of control, *Pcdhg^S/A^* homozygous mutants, and *Pcdhg^CTD^* homozygous mutants using a peptide antibody raised against the final 15 amino acids of the constant domain and a pan-γ-Pcdh antibody (epitope in constant exon 1/2) reveals that *Pcdhg^CTD^* mutants specifically lack the final 15 amino acids of the third constant exon as expected, but still express truncated γ-Pcdh proteins, apparently at a lower level (see Figure 6). GAPDH antibody was used as a loading control for western blots.

To ensure that the *Pcdhg^S/A^* point mutation resulted in a PKC-nonphosphorylatable constant domain, we immunoprecipitated γ-Pcdh proteins from *Pcdhg^S/A^*and control brain lysates using a pan-γ-Pcdh monoclonal antibody (epitope known to be in constant domains 1 or 2, which are unaffected in either of the two new mutant lines; (Lobas et al., 2012)) and western blotted using an antibody specific to phosphorylated serine substrates of PKC (Fig. 2B). Using this phosphoserine antibody, we found that a portion of immunoprecipitated γ-Pcdhs were phosphorylated by PKC in control lysates, while this was essentially abrogated in *Pcdhg^S/A^* lysates as expected (Fig. 2B). To confirm that γ-Pcdh proteins in *Pcdhg^CTD^* mice lacked the C-terminal 15 amino acids of the constant domain, we performed western blots from control, *Pcdhg^S/A^*, and *Pcdhg^CTD^* brain lysates, and probed with a peptide antibody (Keeler et al., 2015) raised against the final 15 amino acids of the constant domain (Fig. 2C). The expected absence of signal in *Pcdhg^CTD^* homozygotes (but its continued presence in both *Pcdhg^S/A^*homozygotes and control mice) confirmed the proper C-terminal deletion. We also probed with the pan-γ-Pcdh monoclonal antibody, which demonstrated that *Pcdhg^CTD^*mice still express intact γ-Pcdh proteins, albeit at a lower level (Fig. 2C).

### Increased exuberance of cortical dendritic arborization due to nonphosphorylatable γ-Pcdhs in *Pcdhg^S/A^* and *Pcdhg^CTD^* mice *in vivo*

In our prior *in vitro* work, we found that the phosphorylation status of the γ-Pcdh constant domain influences dendritic arbor complexity: cultured cortical neurons overexpressing a nonphosphorylatable γ-Pcdh-A3 (but not a wild-type A3 construct) exhibited significantly more complex dendritic arbors (Keeler et al., 2015). Therefore, we hypothesized that the two new mutant mouse lines, neither of which could undergo PKC phosphorylation of the constant domain, would also display increased dendritic arborization in the cortex *in vivo*.

To assess cortical dendrites *in vivo*, *Pcdhg^S/A^* and *Pcdhg^CTD^* mice were crossed with the *Thy1-YFPH* transgenic line (Feng et al., 2000), in which a subset of cortical layer V pyramidal neurons express YFP, providing a means to trace their morphology. Control, *Pcdhg^S/A^*homozygous, and *Pcdhg^CTD^* homozygous neurons were analyzed using three-dimensional Sholl analysis at 6 weeks of age, a young adult timepoint at which we previously have observed arborization defects in a variety of *Pcdhg* null mutants (Garrett et al., 2012; Molumby et al., 2016; Steffen et al., 2023). As predicted, we found that pyramidal neurons in which all γ-Pcdh proteins cannot be phosphorylated in the constant domain by PKC--due either to the S/A mutation or to loss of the PKC site completely due to C-terminal truncation--exhibit significantly increased dendrite arborization (Fig. 3A-D). This overly exuberant dendritic phenotype resulted from an increase in the number of branch points rather than in the average length of each branch (Fig. 3E, F).

**Figure 3.**
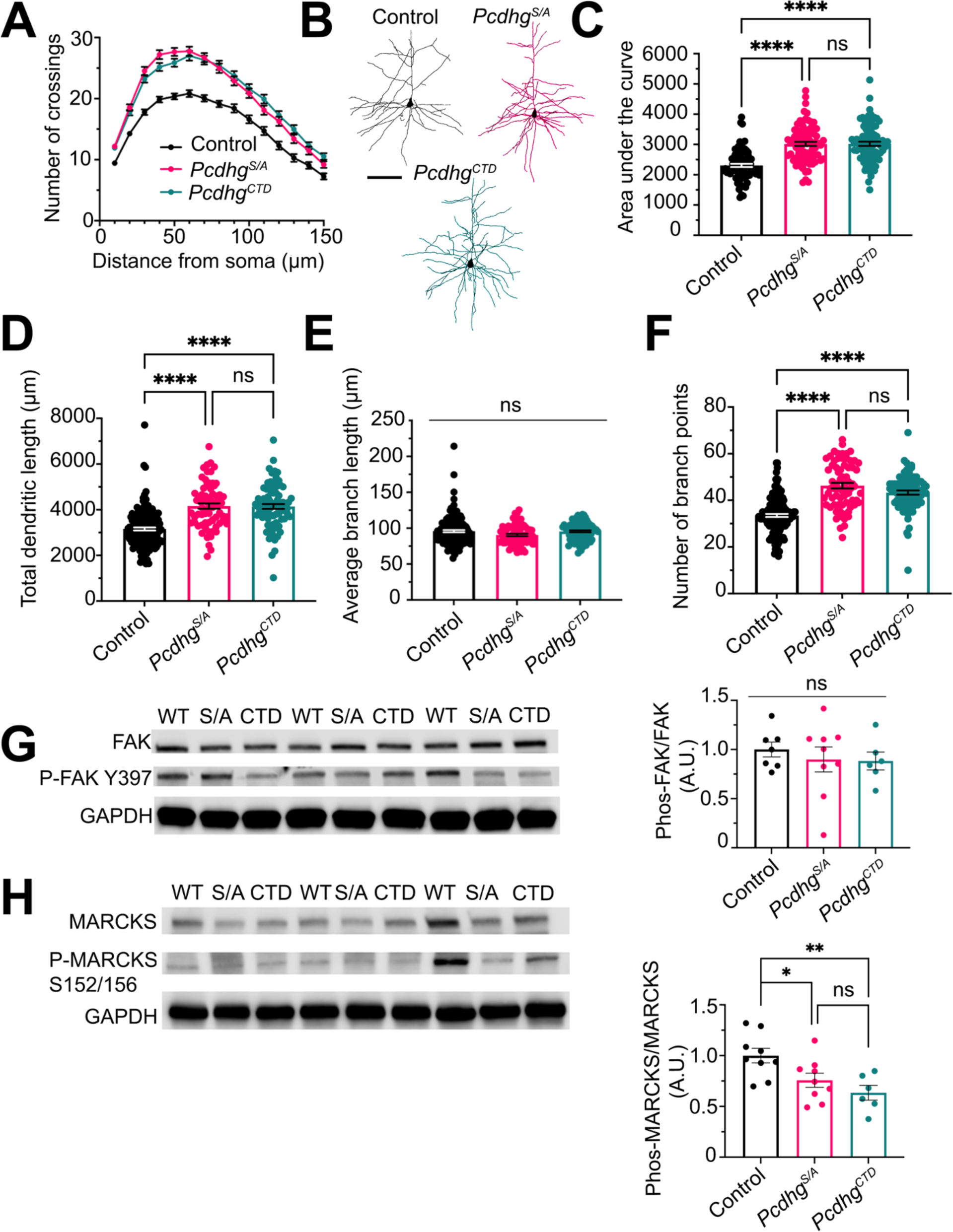
Increased exuberance of cortical dendritic arborization due to nonphosphorylatable γ-Pcdhs in *Pcdhg^S/A^*and *Pcdhg^CTD^* mice *in vivo.* **A,** Sholl analysis graph showing dendritic crossings of concentric circles drawn at increasing 10 μm intervals from the cell body of layer V neurons from control and *Pcdhg^S/A^* and *Pcdhg^CTD^*homozygous mutant mice at 6 weeks of age. **B**, Representative tracings of layer V pyramidal neurons from the three genotypes, color coded as in the graphs. **C**, Graph showing area under the curve of the Sholl graph in panel A. Other graphs indicate total dendritic length (**D**), average branch length (**E**), and number of branch points (**F**). Significant increases in dendritic complexity are seen in *Pcdhg^S/A^*and *Pcdhg^CTD^* homozygous mutant neurons compared to control. N ≤ 80 neurons from 3-4 mice per genotype. Scale bar in **B**: 100 μm. Western blots of P21 brain lysate using pan-FAK and Y397 phosphorylated FAK (P-FAK) antibodies (**G**), or pan-MARCKS and S152/156 phosphorylated MARCKS antibodies (P-MARCKS) (**H**) reveal a non-significant downward trend in relative P-FAK levels in *Pcdhg^S/A^* and *Pcdhg^CTD^* homozygous mutants compared to controls (**G**), and a significant reduction in P-MARCKS levels in *Pcdhg^S/A^* and *Pcdhg^CTD^* homozygous mutants compared to controls (**H**). Duplicate blots were probed with pan- and phospho-specific antibodies, and each were normalized to GAPDH (loading control). Normalized Phos/Pan ratios are plotted as arbitrary units, setting control values at 1. Data are averaged from three replicate experiments. N = 6-9 brains per genotype. One-way ANOVA with Tukey’s multiple comparisons test. Error bars represent SEM; *p<.05, **p<.01, ***p<.001, ****p<.0001; ns = not significant.

No significant differences in arborization were observed between *Pcdhg^S/A^*homozygotes and *Pcdhg^CTD^* homozygotes (Fig. 3A, C-F), suggesting that loss of PKC phosphorylation as a negative regulation point (through amino acid substitution or through loss of the C-terminal motif completely) is the primary reason for the observed over-exuberance in dendritic branching. A potential contribution of the lysine-rich, phospholipid-binding C-terminal motif itself, however, is suggested by the fact that cortical dendrite arborization is significantly increased in heterozygous *Pcdhg^CTD^* mice, but *not* in heterozygous *Pcdhg^S/A^* mice, compared to controls (Fig. 4A-F). Interestingly, this indicates a partially dominant effect of C-terminal motif deletion on arborization, regardless of PKC phosphorylation state.

**Figure 4.**
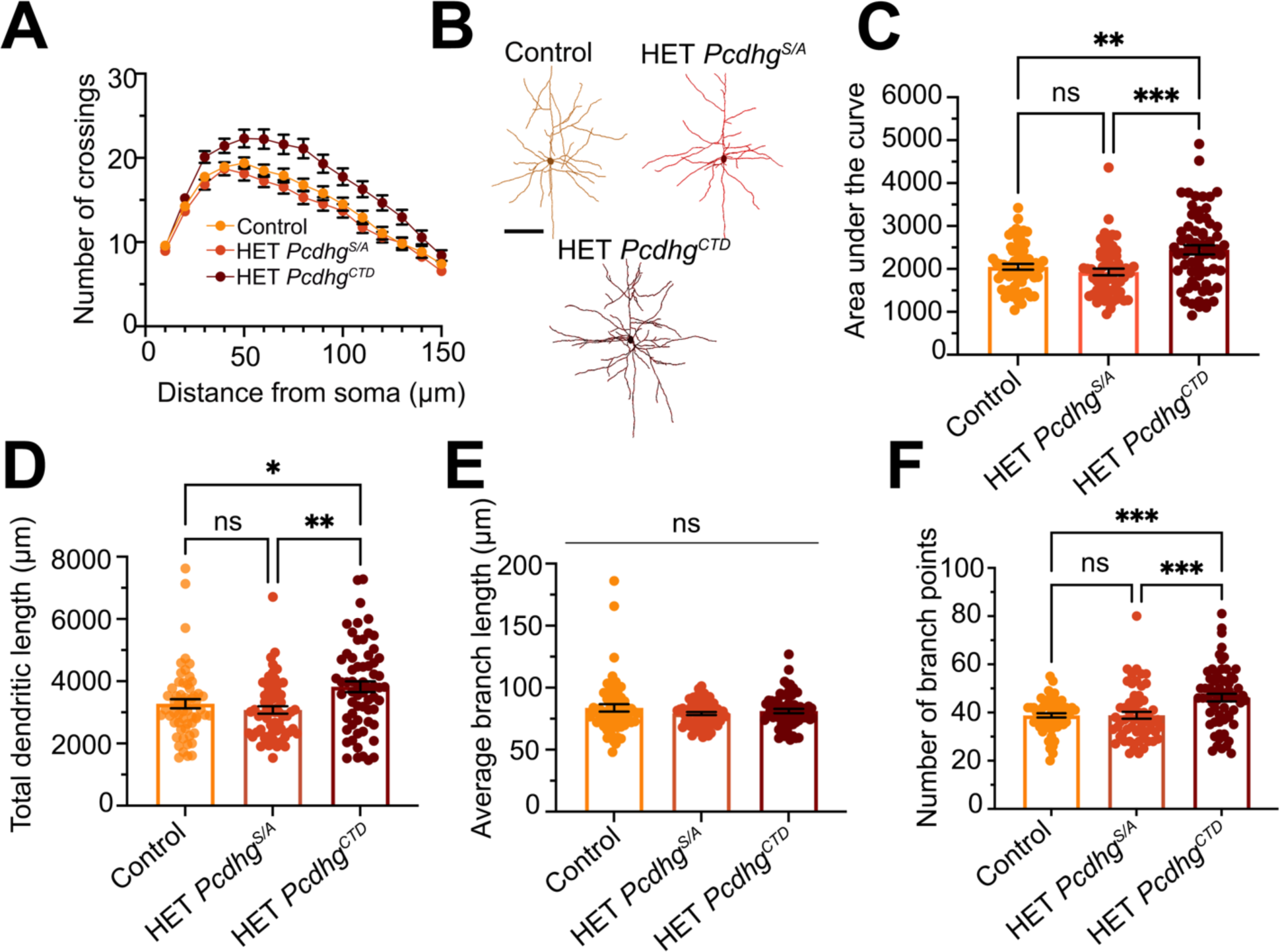
Increased exuberance of cortical dendritic arborization in heterozygous *Pcdhg^CTD^* mice *in vivo*. **A**, Sholl analysis graph showing dendritic crossings of concentric circles drawn at increasing 10 μm intervals from the cell body of layer V neurons from control and heterozygous (HET) *Pcdhg^S/A^* and *Pcdhg^CTD^* mice at 6 weeks of age. **B**, Representative tracings of layer V pyramidal neurons from the three genotypes, color coded as in the graphs**. C**, Graph showing area under the curve of the Sholl graph in **A**. Other graphs indicate total dendritic length (**D**), average branch length (**E**), and number of branch points (**F**). Significant increases in dendritic complexity are seen in heterozygous *Pcdhg^CTD^* compared to controls and heterozygous *Pcdhg^S/A^* mice. Scale bar in **B**: 100 μm. N ≤ 60 neurons from 3-5 mice per genotype. One-way ANOVA with Tukey’s multiple comparisons test. Error bars represent SEM; *p<.05, **p<.01, ***p<.001; ns = not significant.

Our prior work identifying a signaling pathway involving FAK and MARCKS through which γ-Pcdhs influence dendrite arborization (Garrett et al., 2012; Keeler et al., 2015) would predict that the increased dendrite complexity observed in *Pcdhg^S/A^*and *Pcdhg^CTD^* homozygous neurons involves perpetuated inhibition of FAK, a known dendrite arborization inhibitor (Garrett et al., 2012; Rico et al., 2004) as well as the downstream hypophosphorylation of MARCKS, an actin binding protein that promotes dendritic growth (Garrett et al., 2012; Li et al., 2008). We thus asked if we could detect altered signaling in the brains of these new mutant mouse models. Brain lysates from control, *Pcdhg^S/A^*, and *Pcdhg^CTD^* mice at postnatal day 21 (P21; an age at which we previously observed altered signaling in *Pcdhg* null mutants; Garrett et al., 2012) were first subjected to western blotting and probed with a phosphospecific FAK antibody (Fig. 3G). This antibody detects FAK with autophosphorylation at residue Y397, and can thus be used as a readout of FAK activation levels. Duplicate blots were probed with an antibody detecting all FAK, and both FAK and P-FAK levels were normalized to a GAPDH loading control on each respective blot. We did observe a downward trend in FAK activation levels in *Pcdhg^S/A^* and *Pcdhg^CTD^* homozygous brain lysates compared to controls, as expected, but variability between animals prevented this from reaching statistical significance (Fig. 3G).

In contrast, we did observe the expected significant reduction in the levels of MARCKS phosphorylation (Fig. 3H). Hypophosphorylated MARCKS is associated with the plasma membrane and binds to actin. Upon phosphorylation by PKC, MARCKS translocates to the cytoplasm, releases actin, and can no longer promote dendrite complexity (Hartwig et al., 1992; Li et al., 2008; Swierczynski & Blackshear, 1995). Duplicate blots probed with a phosphospecific MARCKS antibody that detects the PKC-targeted serines 152/156 or with an antibody detecting all MARCKS were normalized as described above (Fig. 3H). Statistical analysis revealed a significant decrease in MARCKS phosphorylation levels in both *Pcdhg^S/A^* and *Pcdhg^CTD^* homozygous mice compared to controls. Considering our prior results defining a signaling pathway downstream of the γ-Pcdhs (Garrett et al., 2012; Keeler et al., 2015), this result is consistent with the increased dendritic arborization observed in *Pcdhg^S/A^* and *Pcdhg^CTD^* mice being due, at least in part, to higher levels of active MARCKS.

### Normal density and maturation of dendritic spines in γ-Pcdh C-terminal mutant mice

We found previously that *Pcdhg* null excitatory cortical neurons exhibit increased dendritic spine density, with a preponderance of thin (presumed immature) spines (Molumby et al., 2017). This suggested that γ-Pcdhs negatively regulate the formation of new spines, likely *via* extracellular *cis*-interactions with Neuroligin-1 (Molumby et al., 2017). Given that both FAK (Shi et al., 2009) and MARCKS (Calabrese & Halpain, 2023; Calabrese & Halpain, 2005) have been implicated in the maintenance of dendritic spines, we asked whether γ-Pcdh constant domain signaling might also play a role. To assess dendritic spine density, Vibratome sections of the cortices of control, *Pcdhg^S/A^*homozygous, and *Pcdhg^CTD^* homozygous also harboring the *Thy1-YFPH* allele were prepared, and basal dendritic segments of layer V pyramidal neurons were imaged, and density and spine type analyzed using NeuroLucida software (Fig. 5A; methodology as in Steffen et al., 2023). We found no significant differences between genotypes in either total spine density (Fig. 5B) or in the density of thin, stubby, or mushroom (presumed mature) spines (Fig. 5C).

**Figure 5.**
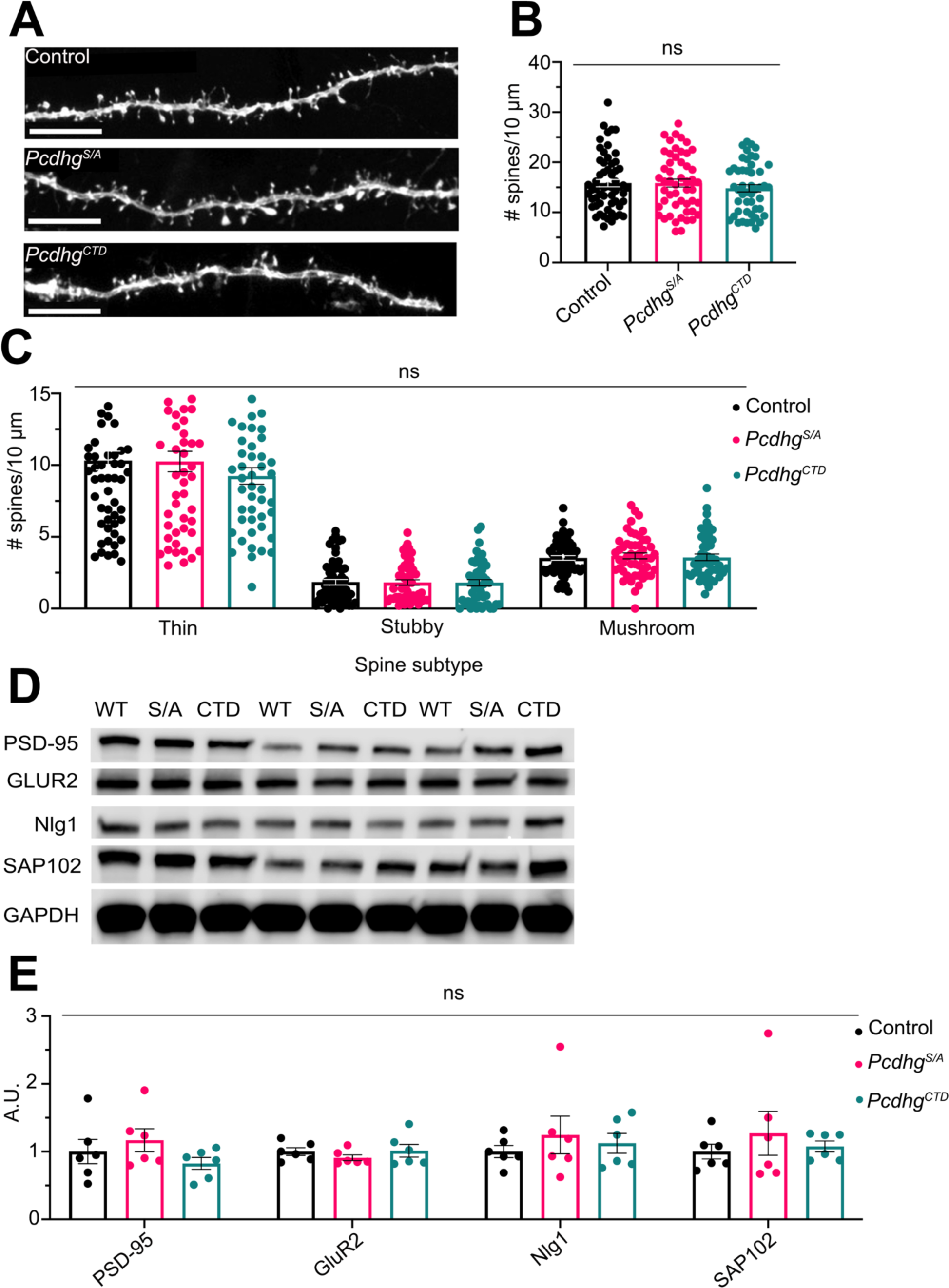
Normal density and maturation of dendritic spines in γ-Pcdh C-terminal mutant mice. **A**, Representative images showing dendritic spines of Thy1-YFPH-labeled layer V pyramidal neurons from the cortex of five-week-old controls and *Pcdhg^S/A^*and *Pcdhg^CTD^* homozygous mutant mice. Scale bar: 5 μm. Quantification of total dendritic spine density (**B**) and of the density of thin, stubby, and mushroom dendritic spines (**C**). No significant differences were observed between controls and either *Pcdhg^S/A^* or *Pcdhg^CTD^* homozygous neurons. N ≤50 dendritic segments from 7-8 mice per genotype. **D**, Western blots of P21 brain lysates reveal no molecular alterations in mutants compared to wildtype (WT) controls, assessed by antibodies against markers of post-synaptic compartments (PSD-95, GluR2, Nlg1, and SAP102). Blots were normalized to GAPDH and values plotted as arbitrary units compared to control values set at 1. Western blotting data were averaged from 3 replicate experiments. N = 6-9 mice per genotype. One-way ANOVA with Tukey’s multiple comparisons test. Error bars represent SEM; ns = not significant.

To seek evidence of any molecular alteration to spines in *Pcdhg^S/A^*and *Pcdhg^CTD^* mutants, we further used western blotting to quantify the levels of proteins localized to, or associated with, post-synaptic compartments. Using a panel of specific antibodies, we quantified levels of PSD-95, GluR2, Nlg1, and SAP102 in control, *Pcdhg^S/A^*, and *Pcdhg^CTD^* P21 brain lysates, normalizing to GAPDH as a loading control (Fig. 5D). We again found no significant differences between genotypes (Fig. 5E), corroborating the lack of alteration in the density and types of dendritic spines. We thus conclude that phosphorylation state and/or presence of the lysine-rich motif of the γ-Pcdh constant domain does not regulate dendritic spine density in the way it does dendritic arborization. This result is not inconsistent with our prior work indicating that γ-Pcdhs primarily affect excitatory and inhibitory synapse density through, respectively, their extracellular interactions with the synaptic organizers Neuroligin-1 and –2 (Molumby et al., 2017; Steffen et al., 2021).

### *Pcdhg^CTD^* mutant brain exhibits significantly decreased γ-Pcdh protein levels

As noted above, qualitative western blot analysis of *Pcdhg^CTD^*mutants revealed an apparent decrease in the levels of γ-Pcdh proteins compared to control (Fig. 2C). This was unexpected, given that the discrete insertion of a single stop codon a mere 45 base pairs prior to the native one would not be expected to activate nonsense-mediated decay or otherwise alter γ-Pcdh transcription or translation. To confirm this reduction of γ-Pcdh protein levels in *Pcdhg^CTD^* mutants, we performed quantitative western blotting of brain lysates to measure levels of all γ-Pcdhs as well as of individual γ-Pcdh isoforms (see Fig. 1A) using a panel of antibodies against: pan-γ-Pcdhs; pan-γ-Pcdh subgroup A isoforms; γB2; γC3; and γC5. Statistical analysis confirmed significant downregulation of the γ-Pcdh protein family overall in *Pcdhg^CTD^* (but not in *Pcdhg^S/A^*) mice, as well as of γ-Pcdh subgroup A isoforms, γB2, and γC5 (γC3 levels trended downward but this did not reach statistical significance) (Fig. 6A, B). The finding that the decrease in overall γ-Pcdh protein levels was not specific to any individual isoform was not surprising, given that the *Pcdhg^CTD^*mutation is in the constant domain, and therefore should affect all isoforms. Overall, these results indicate that discrete C-terminal constant domain mutations can reduce γ-Pcdh protein levels in the brain, which makes the *increased* efficacy of γ-Pcdhs in promoting dendrite complexity in *Pcdhg^S/A^* and *Pcdhg^CTD^* mice all the more remarkable.

**Figure 6.**
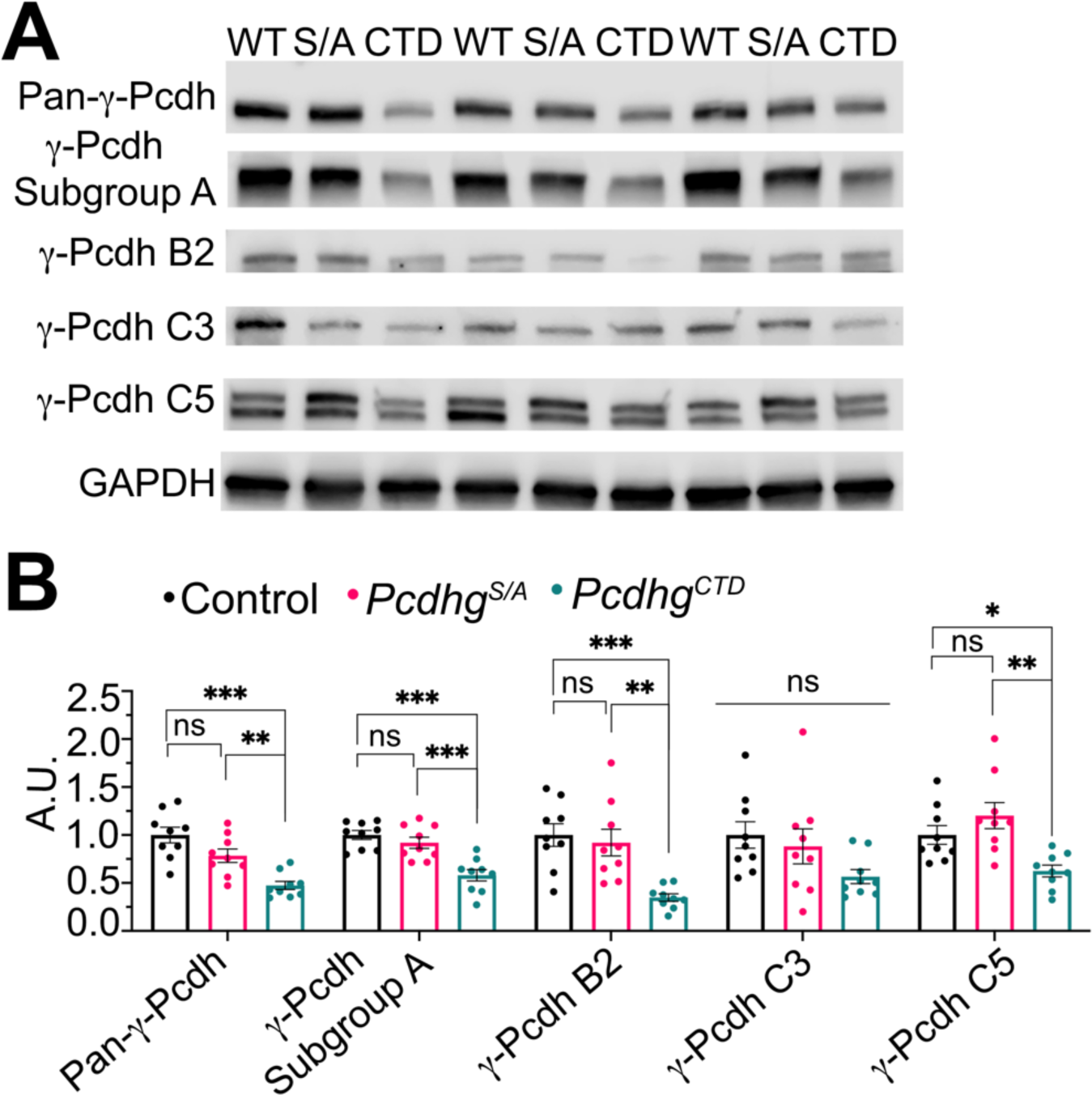
Significantly decreased γ-Pcdh protein levels in *Pcdhg^CTD^* mutant brain. **A**, **B**, Western blots using pan-γ-Pcdh antibody along with a panel of isoform-specific γ-Pcdh antibodies reveal a significant reduction of all γ-Pcdhs, γ-Pcdh subgroup A isoforms, γB2, and γC5 in *Pcdhg^CTD^* homozygous brain lysates compared to controls and *Pcdhg^S/A^* homozygous mutants (a non-significant downward trend in γC3 protein levels was also observed in *Pcdhg^CTD^* mutants). Data were averaged from three replicate experiments. N = 9 mice per genotype. One-way ANOVA with Tukey’s multiple comparisons test. Error bars represent SEM; *p<.05, **p<.01, ***p<.001; ns = not significant.

### Decreased γ-Pcdh protein levels in *Pcdhg^CTD^* neurons are associated with reduced transcript levels and are not attributable to increased protein degradation

To determine if this reduction of γ-Pcdh protein levels in the *Pcdhg^CTD^* brain was a result of protein degradation, we implemented a cycloheximide assay *in vitro*. Cycloheximide inhibits protein synthesis, thus allowing us to track degradation of a discrete γ-Pcdh protein population over time. Day *in vitro* (DIV)8 cultured cortical neurons were treated with 50 µg/mL of cycloheximide for 0 (baseline control), 1, 2, 4, 8, 12, or 24 hours before neurons were lysed and protein samples subjected to western blotting using a pan-γ-Pcdh antibody (Fig. 7A). Levels of γ-Pcdh proteins were normalized to the those at starting time (0h) and plotted for each time point. Statistical analysis revealed no significant differences in γ-Pcdh turnover between control, *Pcdhg^S/A^*, and *Pcdhg^CTD^* neurons (Fig. 7B). These results with cultured neurons *in vitro* suggest that the decreased γ-Pcdh protein levels observed in *Pcdhg^CTD^* brain *in vivo* are unlikely to result from accelerated protein degradation.

**Figure 7.**
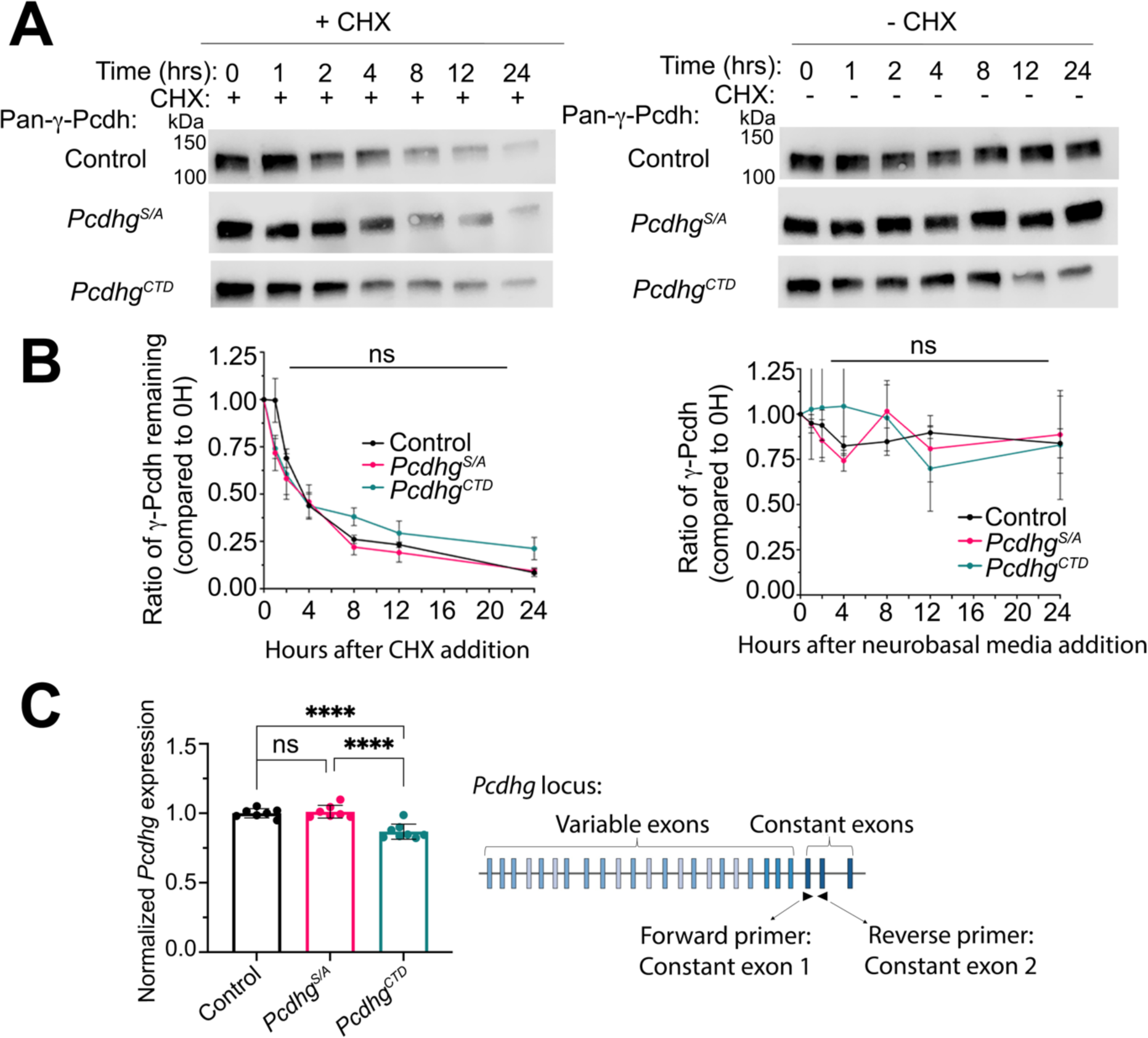
Decreased γ-Pcdh protein levels in *Pcdhg^CTD^*neurons are associated with reduced transcript levels and are not attributable to increased protein degradation. **A,** Western blots of lysates from cultured control, *Pcdhg^S/A^* homozygous mutant, and *Pcdhg^CTD^* homozygous mutant cortical neurons (day *in vitro* (DIV)8) treated with cycloheximide (left) or vehicle (right) for the indicated times and probed with pan-γ-Pcdh antibody. **B**, Ratio of γ-Pcdh protein levels from blots in (**A**) at each time point post cycloheximide addition (or just media change) normalized to baseline levels (0h). Comparison of area under the curves in (**B**) reveals no significant difference in protein degradation rate among genotypes. Data were collected from three separate cultures and experiments. **C**, Quantitative RT-PCR analysis using primers situated in *Pcdhg* constant exon 1 and 2 as shown reveals a significant decrease in pan-*Pcdhg* RNA levels in *Pcdhg^CTD^* homozygous mutant brains compared to controls and *Pcdhg^S/A^* homozygous mutants. Relative *Pcdhg* abundance was calculated using the ΔΔCt method, normalized to GAPDH and control mice. Data were averaged from three replicate experiments. N = 7-8 mice per genotype. One-way ANOVA with Tukey’s multiple comparisons test. Error bars represent SEM; ****p<.0001; ns = not significant.

The next logical step to assessing the decreased γ-Pcdh protein levels in *Pcdhg^CTD^* brains was to investigate *Pcdhg* RNA levels, which we did by performing quantitative RT-PCR. Total RNA was isolated from control, *Pcdhg^S/A^*, and *Pcdhg^CTD^* brains at P21 and first-strand cDNA was synthesized. To quantify total *Pcdhg* transcription, we used a forward primer in constant exon 1 with a reverse primer in constant exon 2 (Fig. 7C). Relative *Pcdhg* expression was quantified using the ΔΔCt method (Livak & Schmittgen, 2001; Winer et al., 1999) to normalize to GAPDH within each sample, and *Pcdhg^S/A^* and *Pcdhg^CTD^* normalized levels were then compared to those of controls. Statistical analysis revealed a significant decrease in *Pcdhg* transcription in *Pcdhg^CTD^* (but not *Pcdhg^S/A^*) brains compared to controls (Fig. 7C). Taken together, these experiments indicate that both *Pcdhg* transcription and resultant γ-Pcdh protein levels are significantly reduced in *Pcdhg^CTD^* mutant brains.

## DISCUSSION

Our prior *in vitro* studies demonstrated that S119 of the γ-Pcdh constant domain shared by all 22 isoforms is a unique target for PKC phosphorylation (Keeler et al., 2015). This serine is embedded in a C-terminal lysine-rich motif that exhibits phospholipid binding *in vitro*, which biochemical studies indicate is abrogated by PKC phosphorylation. PKC phosphorylation of S119 in heterologous cells interferes with the ability of the γ-Pcdh constant domain to inhibit FAK autophosphorylation without affecting its ability to bind to FAK. Overexpression of non-phosphorylatable (S/A mutation) or pseudo-phosphorylated (S/D mutation) versions of one γ-Pcdh isoform (γA3) in cultured neurons suggested that the phosphorylation state of S119 regulates the ability of the γ-Pcdhs to promote dendrite arborization (Keeler et al., 2015). To test whether this mechanism is relevant to neurodevelopment *in vivo*, we report the generation of two new mouse mutants, *Pcdhg^S/A^* and *Pcdhg^CTD^*, in which S119 is rendered incapable of phosphorylation by PKC due, respectively, to an S/A mutation or an early stop codon that creates a 15-amino acid deletion removing the C-terminal motif.

Both mutant lines are viable and fertile and exhibit no gross defects in the architecture of the brain or in cell-type specification in the cerebral cortex, indicating that the unique demonstrated role of one particular γ-Pcdh isoform, γC4, in maintaining neuronal survival (Garrett et al., 2019; Mancia Leon et al., 2024) does not require an intact C-terminus. Consistent with the predictions of our prior *in vitro* results (Keeler et al., 2015), we observe a lack of PKC-phosphorylated γ-Pcdh proteins in the *Pcdhg^S/A^*brain, a significant increase in the dendritic arbor complexity of layer V pyramidal neurons in both *Pcdhg^S/A^* and *Pcdhg^CTD^*, and a corresponding significant decrease in MARCKS phosphorylation in both mutant brains. Additionally, we show no changes in the density of layer V dendritic spines, the proportions of spine morphological types, or the levels of synaptic proteins in *Pcdhg^S/A^* and *Pcdhg^CTD^* brains, consistent with other prior results (Molumby et al., 2017; Steffen et al., 2021, 2023) indicating that γ-Pcdhs regulate synapse development through extracellular *cis*-interactions with Neuroligins rather than through intracellular signaling. Finally, we find a significant reduction in γ-Pcdh protein levels in *Pcdhg^CTD^*, but not *Pcdhg^S/A^*, brains that appears to be due to reduced *Pcdhg* transcription rather than increased protein degradation. Together, these new results confirm and extend several prior observations, improve our understanding of how different γ-Pcdh isoforms and protein domains contribute to the overall functions of the protein family, and suggest several directions for future studies.

While our analysis of *Pcdhg^S/A^* and *Pcdhg^CTD^* mice clearly confirms the importance of the γ-Pcdh constant domain and its C-terminal motif to neural development, it does not yet speak to how constant domain roles may be altered by, or work in tandem with, functions of other γ-Pcdh domains. The γ-Pcdhs form *cis*-dimers (promiscuously among isoforms, though with clear preferences; (Goodman et al., 2022) that engage in strictly homophilic *trans-*interactions (Rubinstein et al., 2015; Schreiner & Weiner, 2010; Thu et al., 2014). Homophilic γ-Pcdh interactions between neurons, and between neurons and astrocytes, have been shown to help drive dendritic arborization in the cortex (Molumby et al., 2016). It remains unknown what impact, if any, the engagement of homophilic interactions has on intracellular signaling *via* the constant domain. It is possible that homophilic binding induces conformational changes in γ-Pcdh proteins that trigger FAK binding/inhibition by the constant domain, or that make S119 less available to be phosphorylated by PKC, allowing for dendrite arbors to grow out. The γ-Pcdh C-terminus is predicted by multiple applications to be “intrinsically disordered”, and we thus do not yet have a good understanding of how protein interactions and phosphorylation events may affect its structure and subsequent function.

A further question raised by the present study is the extent to which signaling roles of particular variable cytoplasmic domains (VCDs; see Fig. 1A) interact with those of the constant domain. We recently showed a role for the γC3 isoform, but not a variety of other isoforms, in promoting cortical dendrite arborization *via* its VCD’s interaction with the scaffolding protein Axin1 (Steffen et al., 2023; see also Meltzer et al., 2023 for a similarly unique γC3 role in spinal cord neurons). Since γC3 VCD binding to Axin1 does not require an adjacent constant domain (Mah et al., 2016), it is possible that the two mechanisms act independently. These questions might be addressed in the future through further genetic manipulation *in vivo*; for example, by coupling restricted cortical knockout of the endogenous *Pcdhg* cluster (such as in Garrett et al., 2012, using *Emx1-Cre* and the conditional *Pcdhg^fcon3^* allele) with attempted rescue of reduced dendrite arborization with: 1) a wild-type γ-Pcdh-C3 isoform (as in Lefebvre et al., 2012); 2) a new line expressing γC3 harboring the C-terminal deletion generated here; or 3) new lines expressing γC3 lacking either the VCD or the constant domain.

Interestingly, we found that the *Pcdhg^CTD^* mutant, which lacks the entire phospholipid-binding motif, appears to have a stronger phenotype than *Pcdhg^S/A^*, which retains the motif: *Pcdhg^CTD^*homozygous mice exhibit somewhat lower (not significantly so, however) levels of phosphorylated MARCKS compared to *Pcdhg^S/A^* homozygotes (Fig. 3H) and *<u>heterozygous</u> Pcdhg^CTD^* neurons have significantly increased dendritic complexity compared to control while *Pcdhg^S/A^* heterozygotes do not (Fig. 4). These effects seem unlikely to be due to reduced γ-Pcdh protein levels in *Pcdhg^CTD^* brains (Fig. 6), since the phenotypes are consistent with *increased*, rather than *reduced*, γ-Pcdh function. This suggests that the phospholipid-binding motif, apart from S119 phosphorylation itself, may play a role in γ-Pcdh functions. While Keeler et al. (2015) found that the C-terminal motif interacts with phosphatidyl inositol phospholipids (but not lysophospholipids or sphingolipids) and that PKC phosphorylation of S119 abrogates this binding, there are no data yet on the functional role this plays. One possible role could be in regulating γ-Pcdh protein trafficking, given that removal of the cytoplasmic domain has been observed to increase cell surface expression, likely due to removal of ER-retention signals (Fernandez-Monreal et al., 2009; Hanson et al., 2010; O’Leary et al., 2011; Schreiner & Weiner, 2010; Thu et al., 2014). Several other studies have shown that a portion of γ-Pcdh protein is associated with membrane vesicles of the endolysosome system, specifically late endosomes (Fernandez-Monreal et al., 2010; Phillips et al., 2003). However, subsequent work demonstrated that retention in these intracellular compartments is mainly mediated by the VCD, not by the constant domain (Fernandez-Monreal et al., 2009; O’Leary et al., 2011; Ptashnik et al., 2023; Shonubi et al., 2015). This would make it unlikely that the stronger phenotype observed in *Pcdhg^CTD^* mice can be attributed to altered protein trafficking. While the *in vivo* function of the C-terminal phospholipid-binding motif remains unknown, our generation of the *Pcdhg^CTD^*line provides an important tool, and our results provide motivation, for pursuing this in future work.

Our *in vivo* analysis of FAK and MARCKS activation in *Pcdhg^S/A^*and *Pcdhg^CTD^* brains provides partial confirmation of the earlier *in vitro* experiments described in Keeler et al. (2015). Based on the γ-Pcdh signaling pathway elucidated by Garrett et al. (2012) and Keeler et al. (2015), we expected to see in both lines a decrease in FAK phosphorylation at Y397 (that is, a decrease in activated FAK) as well as a decrease in MARCKS phosphorylation at S152/S156 (that is, an increase in active MARCKS) accompanying the observed increase in dendritic complexity. While we did observe a downward trend in FAK activation, this was not statistically significant. We did, however, observe a significant decrease in phosphorylated MARCKS, which is at the downstream end of the pathway defined previously (Garrett et al., 2012), and which directly binds actin to induce cytoskeletal changes associated with dendrite arborization. It should be noted that the earlier experiments were quite different: Keeler et al. (2015) overexpressed an HA-tagged WT or S/A-mutant γ-Pcdh isoform in HEK293 cells, treated with PMA to activate PKC, and then immunoprecipitated the FAK associated with the tagged γ-Pcdh. In this assay, they found that PMA resulted in the WT γ-Pcdh-associated FAK, but not the S/A γ-Pcdh-associated FAK, being more active (phosphorylated). While this supported the idea that PKC phosphorylation of γ-Pcdhs at S119 inhibits their ability to abrogate FAK activation, this biochemical assay was performed in heterologous cells *in vitro* using overexpression constructs. The *Pcdhg^S/A^*and *Pcdhg^CTD^* mice go through embryonic and postnatal development with all γ-Pcdhs being non-phosphorylatable, and thus it is possible that homeostatic mechanisms are in place that keep FAK activation within a normal range *in vivo*. In any case, the significant hypophosphorylation of MARCKS observed supports its role downstream of γ-Pcdhs in promoting dendrite arborization, and γ-Pcdhs may activate upstream partners other than FAK to achieve this result.

Additionally, it is important to note that MARCKS has also been implicated in signaling related to dendritic spine maturation and maintenance (Calabrese and Halpain 2005, 2023). Despite the significant increase in active MARCKS in the brains of our new mutants, we observed no significant changes in dendritic spine density or in the distribution of distinct spine morphologies. It will be important in future studies to refine our understanding of how the γ-Pcdh constant domain regulates cell signaling in distinct cellular compartments to affect different aspects of neural development. Knockout of γ-Pcdhs in excitatory forebrain neurons results in increased spine density (primarily of thin spines), and Molumby et al. (2017) presented *in vitro* evidence that this is due to the loss of γ-Pcdh-mediated inhibition of Neuroligin-1, to which γ-Pcdhs bind in *cis via* their extracellular domains. It’s possible that this mechanism—presumably unaffected in *Pcdhg^S/A^* and *Pcdhg^CTD^*mice—is sufficient to compensate for the altered MARCKS activity, and/or that other compensatory mechanisms might be in place to put a brake on dendritic spine density to prevent aberrant synaptic transmission in neurons with an increased total dendritic arbor.

The present study, together with other recent work (Molumby et al., 2017; Garrett et al., 2019; Steffen et al., 2021; Steffen et al., 2023; Meltzer et al., 2023; Mancia Leon et al., 2024), contributes to a converging understanding of γ-Pcdh family function in which distinct roles are played by both individual isoforms and discrete protein domains. Future work aimed at identifying the full panoply of intracellular signaling partners, including both those that work with the shared constant domain and one or more VCDs, will be important to understanding how particular γ-Pcdh functions predominate depending on the neuronal or glial cell type and the stage of neural development.

## METHODS

### Animals

All experiments included male and female mice and were carried out in accordance with the University of Iowa’s Institutional Animal Care and Use Committee and National Institutes of Health guidelines. All mice were healthy and kept under standard housing and husbandry conditions with food and water provided *ad libitum* and 12/12 hour light/dark cycles. All control and mutant mice were on a C57BL/6J background. Control mice for *Pcdhg^S/A^*and *Pcdhg^CTD^* were C57BL/6J wild-type littermates or age-matched C57BL/6J mice from the same colony and housed together. Generation of the novel *Pcdhg^S/A^* and *Pcdhg^CTD^* alleles using CRISPR/Cas9 genome editing is described in the following section. The *Thy1-YFPH* line (Feng et al., 2000) used for dendrite arborization and dendritic spine analysis was obtained from The Jackson Laboratory (https://www.jax.org/strain/003782).

### Generation of *Pcdhg^S/A^* and *Pcdhg^CTD^* alleles

Guide RNA sequences (sgRNA, listed below) were designed to overlap with the desired mutation and analyzed using RGEN tools to maximize efficiency and minimize off target sites. sgRNAs (50 ng/µl) were mixed with *S. pyogenes* Cas9 mRNA (100 ng/µl) and single stranded repair template (sequences below) and microinjected into C57BL/6J zygotes, which were then implanted into pseudopregnant C57BL/6J females. For *Pcdhg^S/A^*, 23 founders were recovered from five females and screened by Sanger sequencing. Of these, 8 harbored indel mutations at the guide site, while one carried the desired mutation, indicative of successful repair from the donor template. For the *Pcdhg^CTD^* allele, 22 founders were recovered from five females, and four had the desired insertion, three of which also had other indels with some mosaicism. Four others were found with indels at the guide site without the desired insertion. For both alleles, founders carrying the desired mutations were bred with C57BL/6J mice and offspring genotyped as described below to screen for germline transmission. Offspring from a single founder per allele were used for the subsequent experiments. Lines were bred out several generations prior to analysis. Mice were genotyped using PCR with the primers indicated in the next section on tail samples, followed by Sanger sequencing to detect the mutations.

#### sgRNA sequences

*Pcdhg^S/A^*: TAATACGACTCACTATAG<u>AATGGCAACAAGAAGAAATC</u>GTTTAAGAGCTATGCTGGAAAC
*Pcdhg^CTD^*: TAATACGACTCACTATAG<u>AAGGCTCCAGCAGGTGGTAA</u>GTTTAAGAGCTATGCTGGAAAC

Protospacer sequence is underlined.

#### Donor sequences

*Pcdhg^S/A^:* CAGAACGTGTACATCCCTGGCAGCAATGCCACGCTGACCAATGCCGCTGGCAAACGAGATGGCAAGGCTCCAGCA GGTGGTAATGGCAACAAGAAGAAA**<u>G</u>**CGGGCAAGAAAGAGAAGAAGTAATATGGAGGCCAGGCCTTGAACCACA AGGCAGCCTCCCTCCCCAGCCAGTCCAGCTCGTCCTTACTTGTACCCAGGCC
*Pcdhg^CTD^:* GCAGCACGTGCCTGACTACCGCCAGAACGTGTACATCCCTGGCAGCAATGCCACGCTGACCAATGCCGCTGGCAA ACGAGATGGCAAGGCTCCAGCA**<u>TAA</u>**GGTGGTAATGGCAACAAGAAGAAATCGGGCAAGAAAGAGAAGAAGTAA TATGGAGGCCAGGCCTTGAACCACAAGGCAGCCTCCCTCCCCAGCCAGTCCA

Sequence divergent from WT is underlined and bold.

### Primers

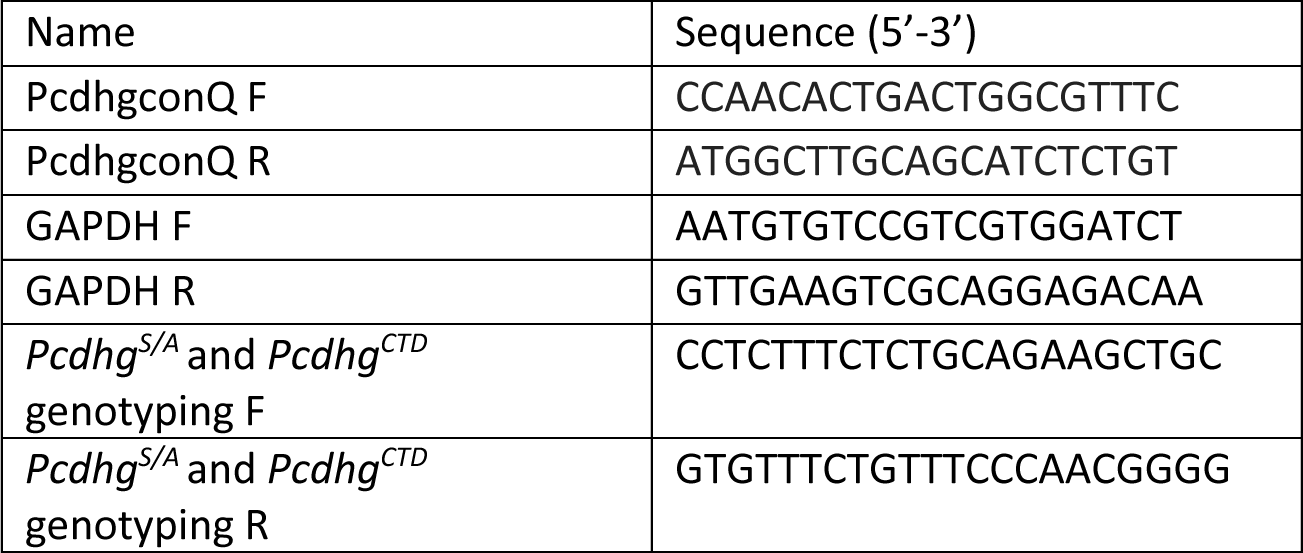

### Quantitative RT-PCR

Whole brains were collected from mice at P21 and RNA isolated using TRIzol reagent (ThermoFisher) according to the manufacturer’s protocol. RNA was further purified using the RNeasy midi cleanup kit (Qiagen) according to the manufacturer’s protocol. Two μg of total RNA was used for cDNA synthesis using High-Capacity cDNA Reverse Transcription kit (ThermoFisher). The cDNA produced was diluted 1:2 for use with LightCycler 480 SYBR Green I Master (Roche) with the primers listed in the section above. The QuantStudio 3 Real Time PCR system (ThermoFisher) was used to carry out qPCR reactions. Relative *Pcdhg* abundance was calculated using the ΔΔCt method, normalized to GAPDH and control mice.

### Tissue preparation and immunofluorescence

Brain tissue from control, *Pcdhg^S/A^*, and *Pcdhg^CTD^* homozygous or heterozygous mice at 6 weeks of age was fixed in 4% paraformaldehyde (PFA) by transcardial perfusion followed by immersion overnight in the same fixative at 4°C. For cryostat sectioning, fixed tissues were sunk in 30% sucrose in PBS at 4°C, immersed in Tissue-Tek OCT compound, and frozen in an isopentane-cooled dry ice/ethanol bath. Cryostat sections were cut at 18 μm and thawed onto gelatin-subbed slides. For immunofluorescence, briefly, sections were rehydrated, blocked with 2.5% BSA, 0.1% Triton X-100 in PBS, and incubated with primary antibody overnight at 4°C in the same solution. The next day, sections were washed with PBS and incubated with fluorophore-conjugated secondary antibody for 1 hour at room temperature (RT). Slides were then washed and DAPI added to the final wash to stain nuclei. FluoroGel (Electron Microscopy Sciences) was used to mount coverslips over the sections.

### Tissue preparation and image collection for dendrite arborization and dendritic spine density analysis

Brains perfusion-fixed as above were embedded in 2% agarose before being coronally Vibratome sectioned at 150 μm (dendritic spines) or 200 μm (dendrite arborization). For dendrite arborization, images were collected from layer V pyramidal neurons in the S1 cortex using a 20x objective on a Leica TCS SPE confocal microscope *via* the Leica LAS X software. Stacks were 100 μm in the Z-dimension with a step size of 0.5 μm. Individual neurons were traced using FIJI’s Simple Neurite Tracer plugin. Sholl analysis was conducted for each individual neuron using FIJI’s Sholl Analysis plugin. Area under the curve for each neuron was calculated using Prism (GraphPad). Researchers were blind to genotype during imaging and analysis.

Dendritic spine density was analyzed as previously described (Anderson et al., 2020; Steffen et al., 2023). Basal dendritic segments (50–200 μm from the soma) of layer V pyramidal neurons in the S1 cortex were imaged using a 100x objective (N.A.= 1.4) on a Leica TCS SPE confocal microscope *via* the Leica LAS X software. Z-stacks were collected with a step size of 0.1 μm. Stacks were deconvolved with Huygens Essential software using the Standard mode of the Express Deconvolution function. Images were saved as 16-bit TIFFs and opened in NeuroLucida 360 software for three-dimensional analysis. Pixel dimensions (0.07 x 0.07 x 0.10 μm) were entered to override the default x and y scaling. Dendritic segments were traced using the Rayburst Crawl user-guided method. To ensure that the tracing encompassed the whole segment, the thickness of the tracing was manually adjusted. Dendritic spines were semi-automatically detected using user-defined parameters (Outer range: 3 μm; minimum height: 0.3 μm; detector sensitivity: 95%; minimum count: 20 voxels). Detected spines were characterized by morphologic subtype using the following parameters: thin or mushroom spines were characterized if the head-to-neck ratio was > 1.1, with spines having a head diameter >0.35 μm classified as mushroom and the remainder considered thin. Spines with a head-to-neck diameter ratio of <1.1 were also classified as thin if the spine length-to-neck diameter ratio was >2.5, with the remainder classified as stubby. Classified spines were then viewed in NeuroLucida Explorer, where spine density was obtained using the Branch Structure analysis. Researchers were blind to genotype during imaging and analysis.

### Cortical neuronal cultures

Cultures were prepared from control (C57BL/6J), *Pcdhg^S/A^*, or *Pcdhg^CTD^* timed matings as described (Garrett et al., 2012) with some modifications. In brief, cortices were isolated from newborn animals and treated with a papain enzyme solution (20 U/ml) for dissociation. The tissues were rinsed briefly in a light inhibitor solution (1 mg/ml trypsin inhibitor, 1 mg/ml BSA) and were then transferred to a heavy inhibitor solution (10mg/ml trypsin inhibitor, 10mg/ml BSA). The tissues were then rinsed 2x in plating media solution (Basal Medium Eagle, 5% FBS, Glutamax, N2 supplements, and 1% penicillin/streptomycin). Cells were triturated in plating media solution and plated onto 12 mm coverslips that had been previously coated with poly-D-lysine and laminin at a density of 250,000-500,000 cells per coverslip. After 6 hours, and then every 2 days subsequent, the medium was changed with fresh Neurobasal Plus (Gibco) supplemented with Glutamax, B27-Plus (Invitrogen) and 1% penicillin/streptomycin (100 U/ml).

### Cycloheximide assay

Primary cortical neurons were cultured as described and plated at a density of 500,000 neurons per 12 mm coverslip. On DIV7, half of the Neurobasal media was replaced with Neurobasal containing 50 µg/mL of cycloheximide (Sigma) or vehicle only. After incubation for 0-24 hours, media was removed and coverslips were briefly rinsed with PBS. Neurons were lysed in RIPA buffer (50 mM Tris-HCl pH 7.4, 5mM NaF, 0.1% SDS, 0.25% sodium deoxycholate, 1% NP-40, 0.15M NaCl) and subjected to SDS/PAGE and Western blotting as described below.

### SDS/PAGE and Western blotting

Brain lysate was prepared by homogenizing forebrains of P21 mice in 1 mL RIPA buffer (50 mM Tris-HCl pH 7.4, 5mM NaF, 0.1% SDS, 0.25% sodium deoxycholate, 1% NP-40, 0.15M NaCl) supplemented with protease inhibitor (Roche miniComplete) and PhosSTOP phosphatase inhibitor (Roche). Protein samples of equal amounts were loaded into TGX precast gels (Bio-Rad), separated via SDS/PAGE, and transferred onto nitrocellulose membranes using a TransBlot Turbo System (Bio-Rad). After protein transfer, membranes were blocked in 5% nonfat milk or bovine serum albumin (BSA, Sigma-Aldrich) in TBS with 0.1% Tween 20 (TBST) for 1 h. Membranes were then washed 3x for 5 min in TBST. Primary antibodies were diluted in 2.5% BSA in TBST and membranes were incubated overnight at 4°C. Blots were washed 3x in TBST. The membrane was then incubated for 1 h in horseradish peroxidase (HRP)-conjugated secondary antibodies diluted in 2.5% BSA in TBST. Signals were detected using SuperSignal West Pico or Femto Enhanced Chemiluminescent Substrates (Thermo Fisher Scientific) on a LI-COR Odyssey FC imager.

### Immunoprecipitation

Whole brain lysates were prepared from control and *Pcdhg^S/A^*mice at 3 months of age. Tissue was homogenized in 2 mL mild lysis buffer (50 mM tris-HCl, pH7.4, 150 mM NaCl, 25 mM NaF, 1% Triton X-100) supplemented with protease inhibitor (Roche miniComplete) and cleared by centrifugation at 1,400 x g for 10 min at 4°C. Protein concentrations were calculated using the Pierce BCA Protein Assay and lysates were diluted to 2 mg/mL. Samples were precleared with 30 μL of Pierce Protein A/G Agarose Beads rotating at 4°C for 1 hour. Samples were immunoprecipitated with 5 μg of pan-γ-Pcdh N159/5 antibody (UC Davis/NIH Neuromab) overnight at 4°C. The next day, 30 μL of beads was added to each tube and rotated for 3 hours at 4°C. Beads were pelleted by centrifugation and rinsed with mild lysis buffer. Beads were resuspended in 50 μL 2x Laemmli buffer, boiled for 10 min, and analyzed by SDS/PAGE and Western blot.

### Experimental design and statistical analysis

Statistical analysis was performed using GraphPad Prism software. Data points on western blotting and quantitative RT-PCR graphs represent individual animals, averaged for each sample from three technical replicates. Data points on dendritic arborization graphs represent individual neurons, and data points on dendritic spine graphs represent individual dendritic segments. Data points on cycloheximide graphs represent individual primary culture experiments. One-way ANOVA with Tukey’s multiple comparisons test was used for comparisons between three groups.

### Antibodies

Primary antibodies are listed below. Secondary antibodies were conjugated with Alexa-488, -568, or - 647 (1:500, Invitrogen) or HRP (1:1000-1:5000, Jackson ImmunoResearch).

| Antibody | Host | Concentration | Company | RRID |
| --- | --- | --- | --- | --- |
| FAK | Rabbit | 1:1000 | Cell Signaling 3285 | AB_2269034 |
| Phospho-FAK Y397 | Rabbit | 1:1000 | Cell Signaling 3283 | AB_2173659 |
| MARCKS | Rabbit | 1:1000 | Thermo Fisher PA5-87527 | AB_2804226 |
| Phos-MARCKS S152/156 | Rabbit | 1:1000 | Cell Signaling 2741 | AB_2140318 |
| Phospho-(Ser)<br>PKC substrate | Rabbit | 1:1000 | Cell Signaling<br>2261 | AB_330310 |
| Pan-γ-Pcdh<br>N159/5 | Mouse | 1:500 | US Davis/NIH<br>NeuroMab 75-185 | AB_2877195 |
| Pan-γ-Pcdh-A<br>N144/32 | Mouse | 1:500 | UC Davis/NIH<br>NeuroMab 75-178 | AB_2877459 |
| γ-Pcdh B2<br>N148/30 | Mouse | 1:500 | UC Davis/NIH<br>NeuroMab 75-184 | AB_2877457 |
| γ-Pcdh C3<br>N174B/27 | Mouse | 1:500 | UC Davis/NIH<br>NeuroMab 75-237 | AB_2877444 |
| γ-Pcdh C5 | Mouse | 1:500 | Invitrogen MA5-26922 | AB_2724958) |
| GAPDH | Mouse | 1:1000 | Abcam 8245 | AB_2107448 |
| γ-Pcdh C-terminal<br>peptide ("GCN") | Rabbit | 1:1000 | From Keeler et al.,<br>(2015) | Available upon<br>request |
| CC3 | Rabbit | 1:100 | Cell Signaling<br>9661 | AB_2341188 |
| GFAP | Mouse | 1:500 | Sigma-Aldrich<br>G3893 | AB_477010 |
| NeuN | Mouse | 1:300 | Millipore MAB377 | AB_2298772 |
| Cux1 | Rabbit | 1:100 | Proteintech<br>11733-1-AP | AB_2086995 |
| Foxp2 | Rabbit | 1:500 | Abcam 16046 | AB_2107107 |
| Parvalbumin | Rabbit | 1:200 | Abcam 11427 | AB_298032 |
| PSD95 | Mouse | 1:1000 | Cell Signaling<br>36233 | AB_2721262 |
| GLUR2 (AMPA2) | Rabbit | 1:1000 | Synaptic Systems<br>182103 | AB_2113732 |
| Nlg1 | Mouse | 1:1000 | Antibodies<br>Incorporated 75-160 | AB_2235964 |
| SAP102 MAGUK<br>scaffold protein<br>N19/2 | Mouse | 1:1000 | Antibodies<br>Incorporated 75-058 | AB_2261666 |

## ACKNOWLEDGEMENTS

This work was supported by grants to J.A.W. and R.W.B.: NIH R21 NS090030 and NIH R01 NS055272

## Notes

### Competing Interest Statement

The authors have declared no competing interest.

